# Integrating GPU support for FreeSurfer with OpenACC

**DOI:** 10.1101/2020.09.03.282210

**Authors:** Jingcheng Shen, Jie Mei, Marcus Walldén, Fumihiko Ino

**Affiliations:** Graduate School of Information Science and Technology, Osaka University, Osaka 565-0871, Japan; Department of Anatomy, Université du Québec à Trois-Rivières, Trois-Rivières, QC G8Z 4M3, Canada

**Keywords:** high performance computing, neuroimaging, medical image analysis, FreeSurfer, OpenACC

## Abstract

FreeSurfer is among the most widely used suites of software for the study of cortical and subcortical brain anatomy. However, analysis using FreeSurfer can be time-consuming and it lacks support for the graphics processing units (GPUs) after the core development team stopped maintaining GPU-accelerated versions due to significant programming cost. As FreeSurfer is a large project with millions of source lines, in this work, we introduce and examine the use of a directive-based framework, OpenACC, in GPU acceleration of FreeSurfer, and we found the OpenACC-based approach significantly reduces programming costs. Moreover, because the overhead incurred by CPU-to-GPU data transfer is the major challenge in delivering GPU-based codes of high performance, we compare two schemes, copy-and-transfer and overlapped-fully-transfer, to reduce such data transfer overhead. Experimental results show that the target function we accelerated with overlapped-fully-transfer scheme ran 2.3× as fast as the original CPU-based function, and the GPU-accelerated program achieved an average speedup of 1.2× compared to the original CPU-based program. These results demonstrate the usefulness and potential of utilizing the proposed OpenACC-based approach to integrate GPU support for FreeSurfer which can be easily extended to other computationally expensive functions and modules of FreeSurfer to achieve further speedup.

## I. Introduction

Magnetic resonance imaging (MRI) is a powerful, extensively used non-invasive imaging technology for creating images of the human body. In clinical practice, it is becoming a standard protocol in the examination and assessment of a variety of neurological disorders and brain injuries [1], e.g., multiple sclerosis [2], hemorrhage [3], traumatic brain injury [4] and brain tumors [5]. MRI has also been increasingly used to study mental illnesses such as major depressive disorder [6], schizophrenia [7], post traumatic stress disorder (PTSD) [8] and obsessive-compulsive disorder (OCD) [9]. Figure 1 shows a sample brain MRI scan from the Alzheimer’s Disease Neuroimaging Initiative (ADNI) database (http://www.adni-info.org), an initiative that collects, validates and utilizes and multi-modal data to understand the progression of Alzheimer’s Disease (AD).

**Fig. 1.**
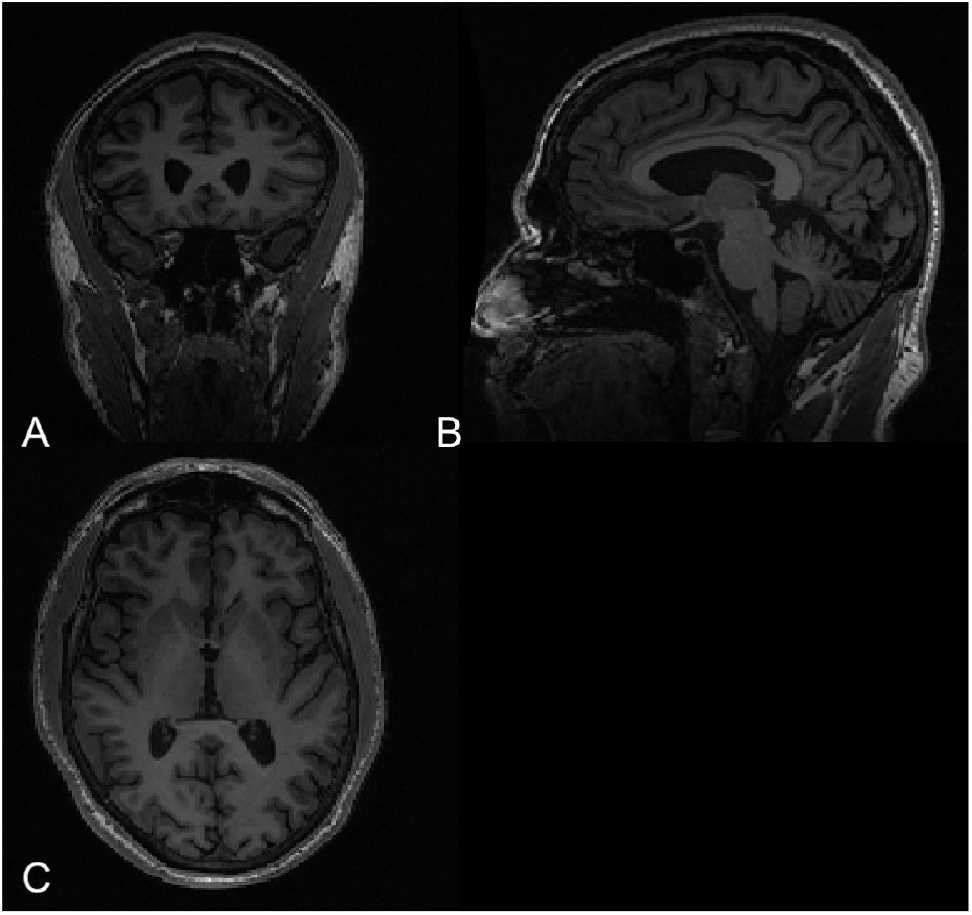
A sample MRI scan from ADNI database displayed from (A) coronal plane, (B) sagittal plane and (C) axial plane.

FreeSurfer [10] is a suite of software tools developed for the study of cortical and subcortical brain anatomy and is widely used by researchers in many subfields of neuroscience and neurology. It allows for quantification of structural, functional and connectional properties of the human brain through analysis of neuroimaging data, i.e., T1-weighted MRI scans. FreeSurfer runs on multiple software platforms and operating systems, and has been made freely available through GitHub [11], enabling parallelization and other modifications to improve its ease of use. Despite the standardized and automated analysis pipeline of FreeSurfer, its execution time can be up to more than 10 hours for analysis of a single MRI scan with a single CPU thread. Although FreeSurfer uses CPU multi-threading with OpenMP [12] to reduce its execution time, it will still consume 2–3 hours even with tens of threads on a manycore CPU.

GPUs are more powerful accelerator devices than manycore CPUs for computing- and memory-intensive applications [13]–[16]. CUDA [17] is a parallel computing platform based on C++ which can be used to access the instruction set and computational elements on Nvidia GPUs. Given that FreeSurfer is a large-scale project with more than 100 million lines of source code and is being updated frequently to add new features and bug fixes, to create a CUDA-based GPU acceleration of FreeSurfer, developers must rewrite the computation kernels with CUDA to allow compatibility with code changes and version updates of FreeSurfer. As a result of the huge programming costs, the core development team of FreeSurfer stopped integrating the CUDA-based GPU acceleration into FreeSurfer after the release of FreeSurfer 5.0 in 2012 [18].

To reduce FreeSurfer’s execution time, in this study we propose a novel yet realizable approach to integrate GPU acceleration into the latest 7.1.0 version of FreeSurfer with a reasonable programming cost using OpenACC [19], a directive-based programming paradigm similar to OpenMP. Compared to CUDA-based GPU acceleration, OpenACC requires almost no modification to the source codes except insertions of directives into the code regions to parallelize for the acceleration of the programs. The OpenACC directives instruct the compilers (e.g., the PGI compiler) to parallelize the code and offload the work to the GPU.

The primary contributions of this work are summarized as follows:

- We accelerate the most time-demanding module of the latest version of FreeSurfer (i.e., `mri_ca_register`) through GPU parallelization of the function `gcamComputeSSE()` with an efficient scheme to reduce data transfer overhead.
- We introduce a novel, cost-efficient approach to further accelerate FreeSurfer and propose future improvements to OpenACC for porting computation-heavy tasks.

The remainder of this study is organized as follows. Previous studies that accelerated FreeSurfer using high performance computing (HPC) techniques are introduced in Section II. How the OpenACC framework could be used to parallelize CPU-based code is briefly described in Section III. The selection of target kernels to accelerate in this work is explained in Section IV. The proposed method used to parallelize the CPU-based kernels is described in Section V. In Section VI, experimental results are presented and analysed. Finally, Section VII concludes the present study and suggests future research directions.

## II. Previous Work

Pantoja *et al.* [18] proposed a hybrid approach to accelerate FreeSurfer with both a GPU and a multicore CPU. In their study, CPU-based code runs with OpenMP threads, which is the same as the original implementation of FreeSurfer. Moreover, they used CUDA to rewrite a kernel for GPU acceleration. However, this study has two major limitations. First, although not specified, the authors appear to have worked on the obsolete FreeSurfer version 5.0, which was released in 2012. Secondly, the kernel that was rewritten with CUDA in this work is no more a dominant part of the latest version of FreeSurfer, therefore the acceleration achieved will be relatively limited if their method is applied to the latest version.

Delgado *et al.* [20] proposed a pipeline scheme to schedule the execution of modules of FreeSurfer, achieving a task-level parallelism. For the GPU acceleration, they used the last version of FreeSurfer that supports GPU, i.e., the now obsolete FreeSurfer version 5.0.

Giuliano *et al.*’s work [21] showed that running FreeSurfer on a hybrid CPU-to-GPU infrastructure can lead to a reduced execution time compared to using a CPU-only infrastructure. Nevertheless, same as in [18] and [20], an obsolete version of FreeSurfer was used.

Madhyastha *et al.* [22] and Zablah *et al.* [23] explored the possibility of running FreeSurfer on cloud computing services and evaluated the associated performance. In [22], the authors investigated the benefits of using Amazon Web Services (AWS) to run neuroimaging applications including FreeSurfer, and found that no within-brain (i.e., MRI scan) parallelism occurred when an MRI scan is processed on a single workstation with FreeSurfer. Moreover, they did not conduct experiments using GPU-accelerated FreeSurfer because no GPU-based version was available except for the obsolete FreeSurfer 5.0. In [23], the authors evaluated the feasibility of using FreeSurfer on cloud computing environments by running FreeSurfer on virtualized cloud infrastructures and physical platforms, then compared the performance. They found the penalty for the use of virtualized infrastructures was less than 4.5%, concluding that FreeSurfer can benefit from the scalability of cloud services that outweighs the small overhead incurred by using cloud services. GPU acceleration for FreeSurfer is not mentioned in [23].

Given the limitations of previous works, in the present study, we accelerate three kernels including one kernel accelerated in [18]. We utilize OpenACC directives to generate GPU-based code, which significantly reduces programming costs caused by manually rewriting kernels with CUDA. We also present a overlapping scheme with inter-operating OpenACC directives using OpenACC and CUDA APIs, which hides the data transfer overhead while requiring minimal modification of existing codes. We believe our approach is more sustainable in terms of programming costs, ease of implementation and flexibility, especially when applied to large scale software suites.

Integrating our OpenACC-based GPU implementation into the task scheduler proposed in Delgado *et al.* [20] should be beneficial for the speedup of current and future versions of FreeSurfer. Additionally, a GPU-accelerated implementation of FreeSurfer can achieve more speedup in a cloud environment compared to a CPU-only implementations such as those used in [22] and [23].

## III. Openacc Framework

OpenACC is a directive-based programming standard for delivering parallel computing programs on heterogeneous computing systems such as manycore CPUs and GPUs. In contrast to CUDA, OpenACC only requires minimal modifications of the CPU-based codes to support GPU acceleration. OpenACC users only need to insert directives into the code regions they want to parallelize, as these directives will instruct the compiler to generate GPU-based codes. Moreover, OpenACC also provides a variety of APIs that allow its users to have more explicit and flexible control over the details concerning parallelization.

Briefly, OpenACC directives are designed to parallelize affine loops. Figure 2 shows a simplified description of one kernel we parallelized in this work with OpenACC directives. This segment of code sums up all the `label_dist` values of valid nodes of the volume. Line 4 copies all the nodes to the GPU for processing. Line 5 instructs the compiler that a nested affine loop of three levels (`loop collapse(3)`) will follow, in which the variable sse will be summed up (`reduction(+:sse)`) and all necessary data is already on the GPU (`present(h_nodes)`). Line 17 deletes the on-GPU data to free the space once the data is not needed anymore. As GPU support could be added to CPU-based codes at low programming costs using OpenACC, the framework is very suitable for porting large existing projects such as FreeSurfer.

**Fig. 2.**
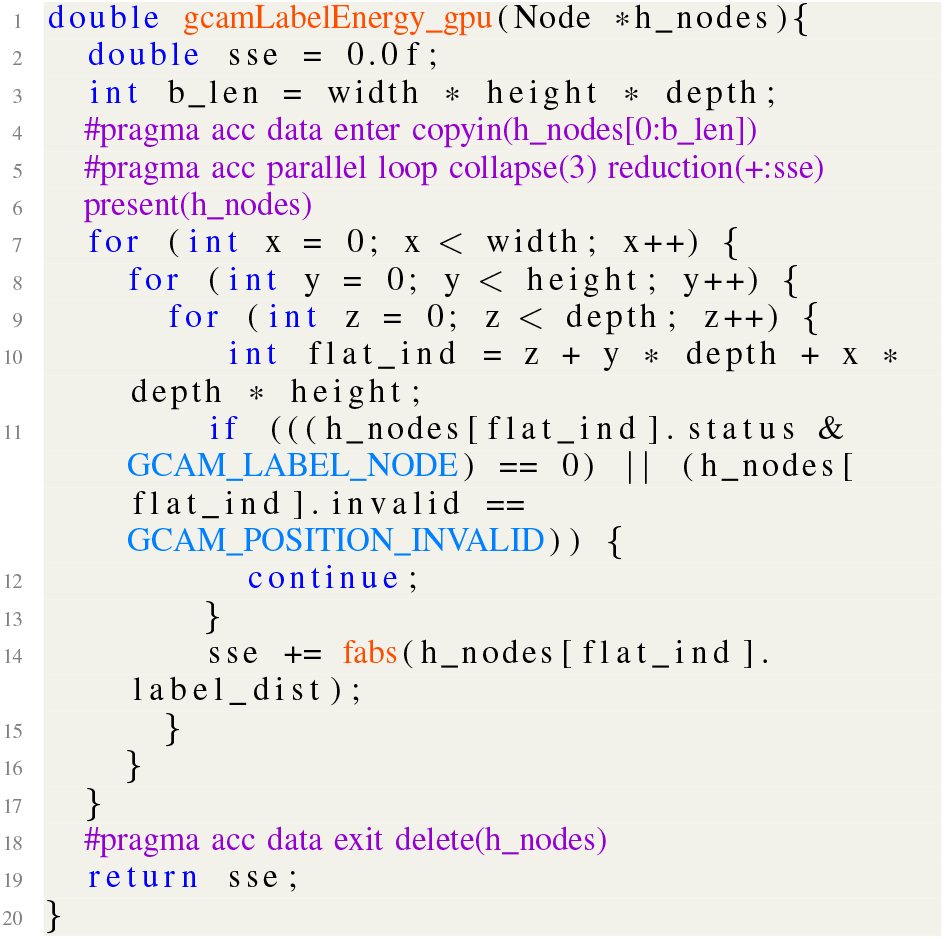
Implementation of the GPU-based version of a CPU-based kernel using OpenACC directives.

## IV. Target Kernels for Acceleration

FreeSurfer comprises multiple component modules (i.e., programs) that are called by the MRI processing stream. To find the target module(s) to accelerate, we applied a standard processing stream to one randomly selected MRI scan from the ADNI database. The dimension of this scan was 256 * 256 * 256. Figure 3 illustrates the execution time of each module involved in the processing stream. The most time-consuming module is `mri_ca_register` (shown in orange) which takes approximately 50 minutes to execute with 32 CPU threads. Therefore, in this work, we aim to accelerate `mri_ca_register` on the GPU by applying OpenACC directives to a function named `gcamComputeSSE()`, which is the most frequently called function by the `mri_ca_register` module. This function is also one of the most compute-intensive functions.

**Fig. 3.**
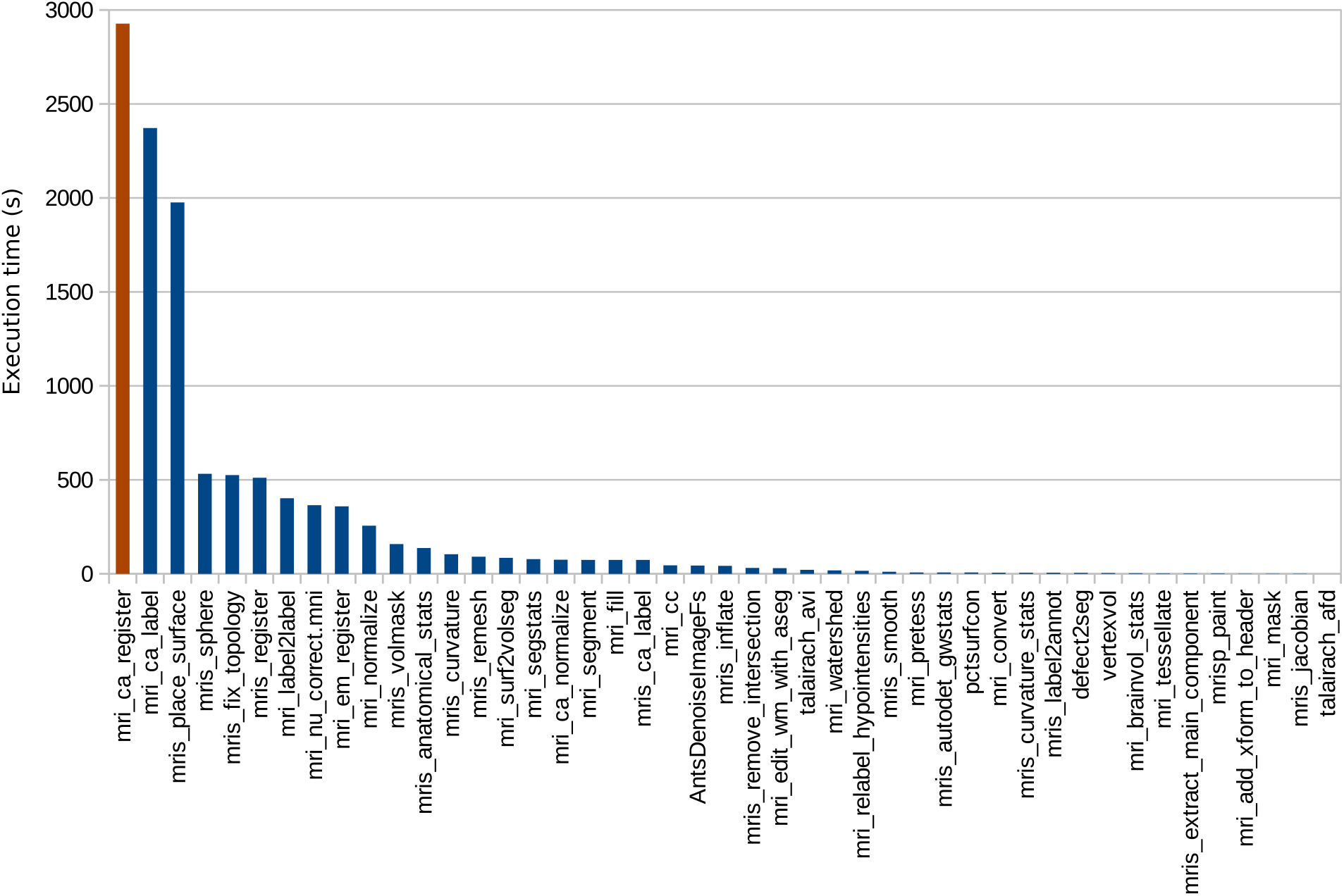
Breakdown of the execution time of a standard FreeSurfer processing stream and a comparison of the individual execution times of its 32 threads.

The function `gcamComputeSSE()` has four computation kernels: `gcamLogLikelihoodEnergy(), gcamLabelEnergy(), gcamSmoothnessEnergy()`, and `gcamJacobianEnergy()`. Although previous work [18] has rewritten the `gcamSmoothnessEnergy()` kernel with CUDA, for minimized programming costs, we accelerate the same kernel with OpenACC directives. In addition to `gcamSmoothnessEnergy()`, we also implement GPU-accelerated versions of two other kernels: `gcamLabelEnergy()` and `gcamJacobianEnergy()`.

## V. Proposed Method

In this work, we aim to accelerate the function `gcamComputeSSE()`. This function accepts an array of nodes as the parameter and computes a sum-of-squared-error that sums up the output of the four kernels `gcamLogLikelihoodEnergy(), gcamLabelEnergy(), gcamSmoothnessEnergy()`, and `gcamJacobianEnergy()`. The nodes are of a composite type with 40 member variables (17 `double`, 15 `float`, 6 `int`, one `char`, and one pointer to another data structure). Note that the four kernels only reference the values of the nodes and do not modify these values at runtime. Two major challenges of accelerating `gcamComputeSSE()` are:

- Each kernel only references less than ten member variables of the node structure; however, we cannot partially transfer the array of structures (AoS) for each kernel as only continuous data transfer is allowed. Therefore, we must reduce the effect of the unnecessary data transfer.
- The first kernel `gcamLogLikelihoodEnergy()` accesses noncontinuous data, i.e., the pointer member of the node structure. Data connected by pointers cannot be easily transferred with the AoS to the GPU because they are noncontinuous.

To overcome the first challenge, we compared two schemes, *copy-and-transfer* and *overlapped-fully-transfer*, that reduce the side effects incurred by data transfer. As for the second challenge, we leave the first kernel `gcamLogLikelihoodEnergy()` on the CPU side to be executed by multiple CPU thread, and this strategy is considered the key to our proposed mapping scheme. Also note that we did not use tiling techniques to overlap data transfer with GPU computations because (1) dividing the data into chunks and tuning the size of these chunks significantly increase the programming costs, and (2) in this study, data transfer overhead, which is estimated to be of approximately tens of ms, cannot be hidden by being overlapped with the GPU computing time, which is normally less than 0.1 ms per call to a kernel.

### A. Copy-and-Transfer

In this scheme, we aim to reduce the data transfer of the array of nodes. We copy only the necessary member variables to buffers and then transfer the buffers to the GPU. By doing so, we in effect partially transfer the AoS at the expense of CPU execution time that increases due to copying from the original AoS to the buffers. Moreover, the same member variables used by different kernels are copied and transferred only once. Figure 4 shows a simplified description of this scheme.

**Fig. 4.**
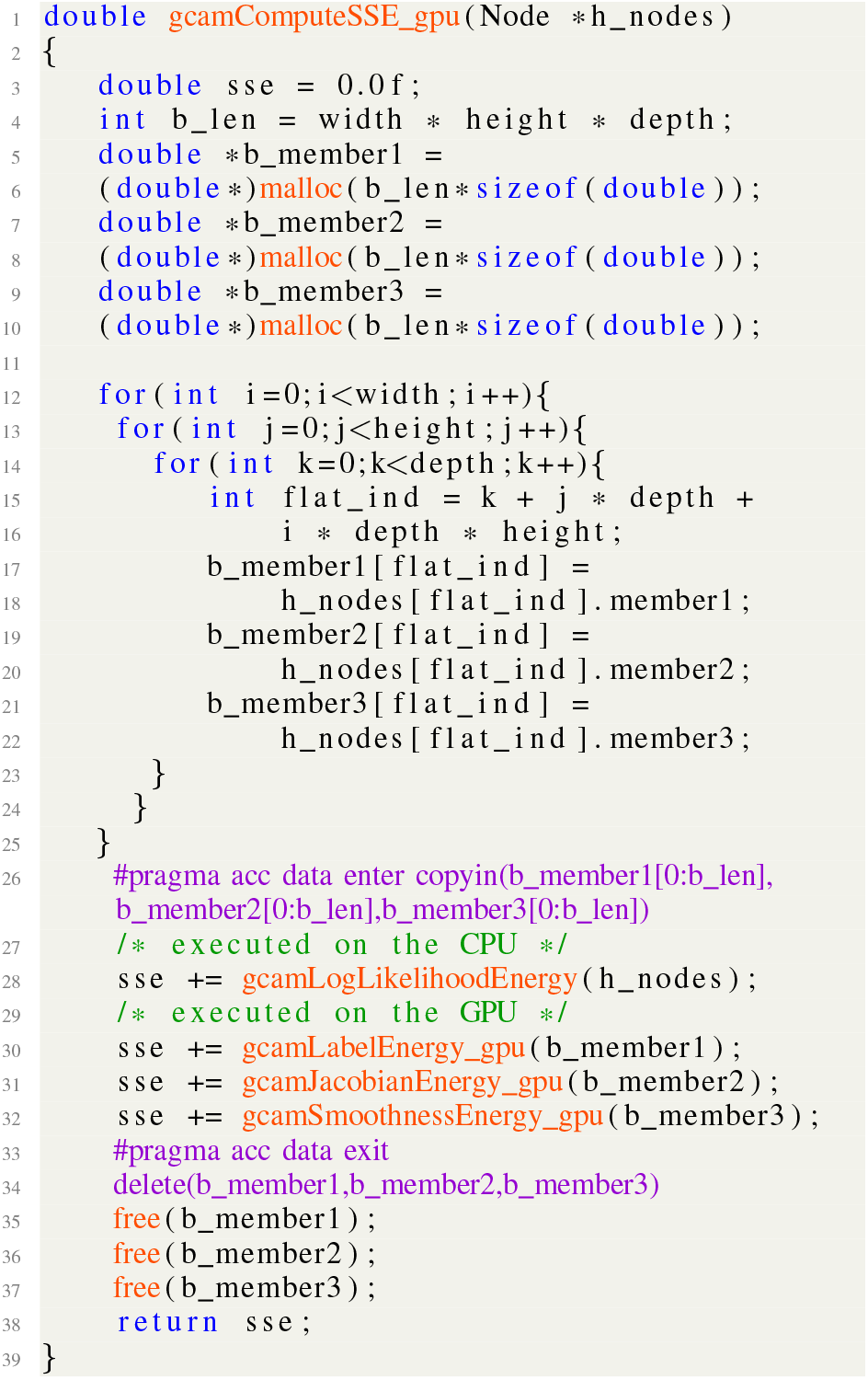
Simplified description of the copy-and-transfer scheme. We copy the necessary member varibles of the nodes to buffers and then transfer the buffers to the GPU with OpenACC `data` directives. OpenACC `parallel loop` directives are applied to the affine loops in the kernels (like Line 5 in Fig. 2).

Since the previous work [18] only accelerated the kernel `gcamSmoothnessEnergy()`, in this scheme, we implement both one-kernel and three-kernel versions to determine how much speedup we can achieve by accelerating two more kernels.

### B. Overlapped-Fully-Transfer

In this scheme, instead of partially transferring the AoS by copying necessary member variables from the original data to buffers, we fully transfer the AoS to the GPU except the data connected by pointers. The overhead of data transfer is relatively high because each node has 40 member variables, summing up to 250 byte per node and therefore, approximately 500 MB in total for about two million nodes. In order to reduce the data transfer overhead, in the present study, the data transfer and the execution of the first kernel `gcamLogLikelihoodEnergy()` on the CPU are made to occur in parallel. Such a procedure is possible because the kernels only reference the member variables without modifying, and can thus be executed simultaneously with CPU-to-GPU data transfer. Figure 5 shows a simplified description of this scheme.

**Fig. 5.**
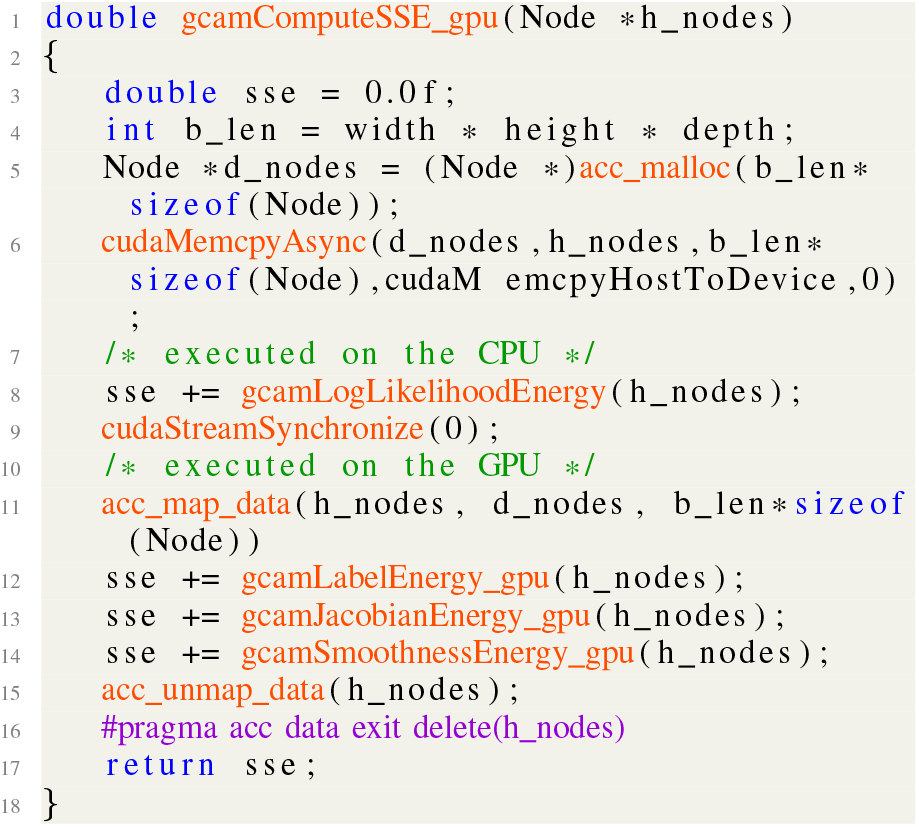
Simplified description of the overlapped-fully-transfer scheme. We interoperate OpenACC directives with OpenACC and CUDA APIs, allowing explicit control of the on-GPU data. OpenACC `parallel loop` directives are applied to the affine loops in the kernels.

In practice, apart from inserting OpenACC directives to code regions to be parallelized, additional efforts are required to realize this scheme. Subject to current NVIDIA GPU driver software, the overlapping of CPU-GPU data transfer with CPU or GPU computations necessitates the use of page-locked (i.e., pinned) memory. However, OpenACC only provides a compiler option `−ta=tesla:pinned` to enable the use of pinned memory. Such a compiler option replaces `malloc()/new` calls with calls to `cudaMallocManaged()` (and `free()/delete` with `cudaFree()`). For a project with thousands of files, such a policy is not desired because the allocation of pinned memory requires more time than the allocation of normal memory, and not all the functions (such as CPU-based ones) benefit from pinned memory. Accordingly, instead of compiling the project with `−ta=tesla:pinned`, we explicitly allocate the array of nodes as pinned memory using a CUDA API `cudaMallocHost()`. Moreover, because the nodes are allocated outside `gcamComputeSSE()`, we utilize a CUDA API (i.e., `cudaMemcpyAsync()`) to asynchronously transfer the nodes to the GPU, explicitly indicating the amount of data to be transferred. OpenACC APIs `acc_map_data()` and `acc_unmap_data()` are used to indicate where are the data for processing on the GPU.

## VI. Experimental Results

To measure the execution time, we applied `mri_ca_register` to MRI scans with size of 256 * 256 * 256 obtained from the ADNI database (http://www.adni-info.org). The processing times of six scans are evaluated in this section. Table I shows the testbed for all experiments performed.

**TABLE I:** Testbed for experiments

|  |  |
| --- | --- |
| GPU | NVIDIA Tesla V100-PCIe |
| CPU | Xeon Silver 4110 |
| OS | Ubuntu 16.04.6 |
| FreeSurfer | 7.1.0 |
| PGI compiler | 19.10 |

### A. Evaluation of Function-wise Speedup

We evaluate the speedups achieved by the proposed method in terms of the execution time of `gcamComputeSSE()`. Four implementations of `gcamComputeSSE()` were considered in the experiment. The first is the baseline, i.e., original CPU-based code that runs with 32 CPU threads. The second implements only one GPU-based kernel, i.e., `gcamSmoothnessEnergy()` using the copy-and-transfer scheme, mimicking the behavior of the previous work [18]. The third and the fourth implement three kernels using copy-and-transfer and overlapped-fully-transfer schemes, respectively. Figure 6 illustrates the exection times of the four implemetations. The execution times of GPU-accelerated kernels (i.e., `gcamLabelEnergy()` (light blue bars), `gcamJacobianEnergy()` (dark red bars), and `gcamSmoothnessEnergy()` (light green bars)) are negligible compared to that of CPU-based kernels. The overlapped-fully-transfer scheme (72 ms) performed the best in the four implementations, running 2.3× as fast as the CPU-based implementation (167 ms). The copy-and-transfer scheme achieved the speedup because the execution time of `gcamLogLikelihoodEnergy()` on the CPU side (about 33 ms) hid two thirds of the CPU-to-GPU data transfer time (about 45 ms); therefore, we can benefit from running `gcamLabelEnergy()`, `gcamJacobianEnergy()`, and `gcamSmoothnessEnergy()` on the GPU only at a small cost (i.e., the unhidden transfer time which is about 11 ms). The copy-and-transfer scheme ran slower than the overlapped-fully-transfer scheme as it involves a buffer copying time (about 27 ms) and a CPU-to-GPU transfer time (about 19 ms). Note that CPU-to-GPU transfer time in the copy-and-transfer scheme was not overlapped because the buffers were allocated as normal memory instead of pinned memory. Buffers are local variables of the kernels and thus are allocated every time the kernels are called. Therefore, allocating buffers as pinned memory per call will result in a significant overhead. Using static variables as buffers to reuse allocated pinned memory may be a solution, but increases the risk of memory leaking because it is difficult to determine when to free the pinned memory. Nevertheless, even if we manage to hide the buffer transfer time, the CPU buffer copying time (27 ms) is still about thrice the overhead of the overlapped-fully-transfer scheme (11 ms). We however notice that CPU-based kernel `gcamLogLikelihoodEnergy()` spent more time in the overlapped-fully-transfer scheme than in the other implementations. This is because overlapping CPU-to-GPU data transfer with CPU computation involves competition in reading from the original data. This side effect may be eliminated by GPU hardware improvement in the future.

**Fig. 6.**
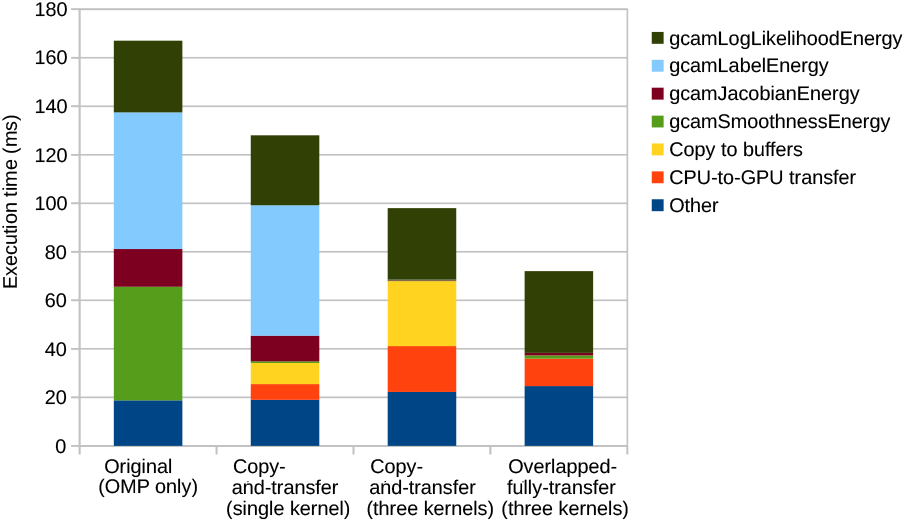
Breakdown of the execution time per call of the four implementations of `gcamComputeSSE()`. Bars with different colors represent the kernel computation time, buffer copy time, CPU-to-GPU data transfer time, and other operation time. The original implementation was executed by only CPU (i.e., OpenMP) threads. Copy- and-transfer (single-kernel version) accelerates `gcamSmoothnessEnergy()` on a GPU, mimicking the behavior of [18]. Copy-and-transfer and overlapped-fully-transfer (three-kernel version) accelerate `gcamSmoothnessEnergy()`, `gcamJacobianEnergy()`, and `gcamLabelEnergy()` on a GPU.

### B. Evaluation of Module-wise Speedup

We evaluate the speedups achieved by the proposed method in terms of execution time of the entire `mri_ca_register` module. The same four implementations described in Section VI-A were used in the experiment. Figure 7 illustrates the execution times of the four implementations. With GPU acceleration and efficient hiding of the CPU-to-GPU transfer overhead, the implementation with overlapped-fully-transfer scheme ran 1.17 as fast as the original CPU-based implementation on average. The highest speedup, 1.23, was achieved by processing the scan (e), benefiting from the largest number of calls to `gcamComputeSSE()` (4869 times). Further speedup can be easily achieved by applying our approach to other functions (blue bars), which is rather repetitive work than a technical contribution and thus not considered in this work.

**Fig. 7.**
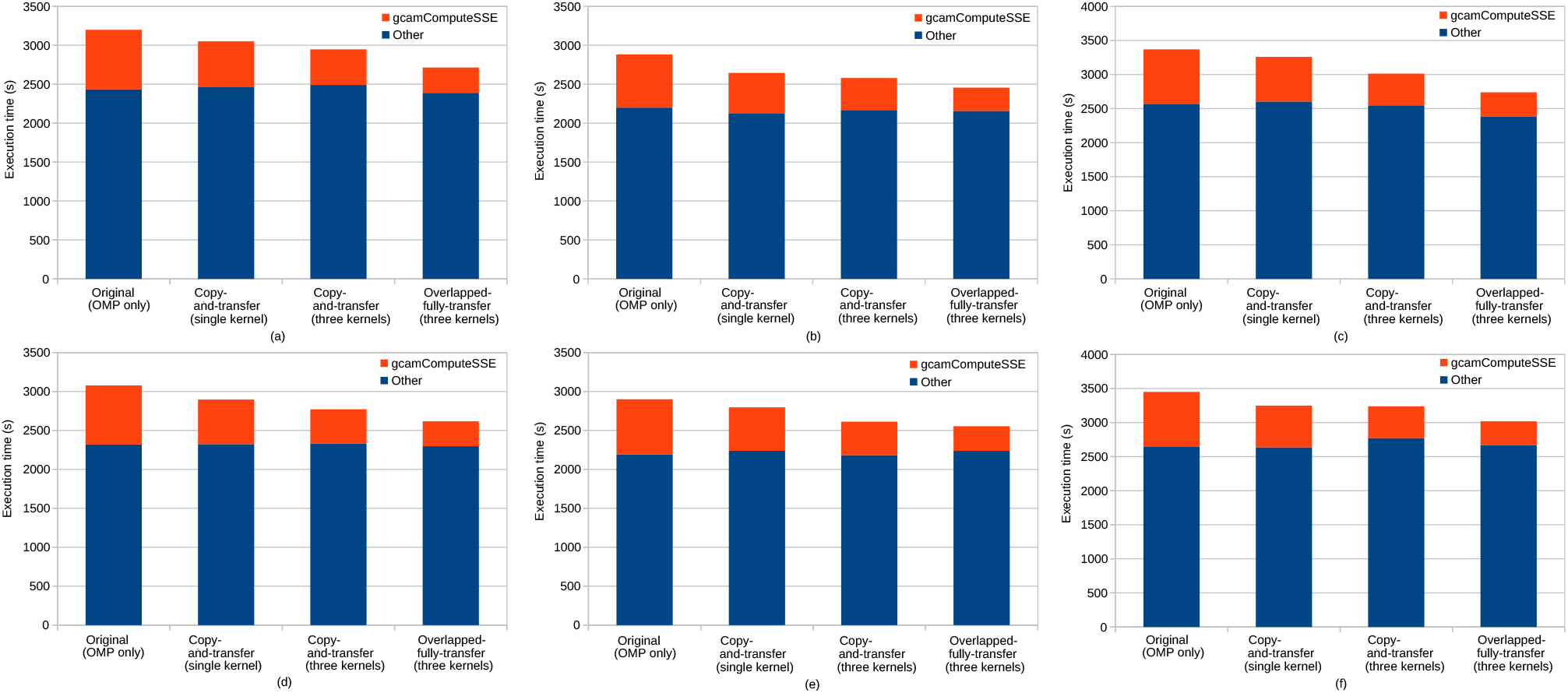
Breakdown of the execution time of `mri_ca_register` implemented with four versions of `gcamComputeSSE()`. Sub-figures (a) to (e) show the execution time when applying the four implementations to the six randomly selected MRI scans, respectively. Note that the number of calls to `gcamComputeSSE()` varies when `mri_ca_register` processes different scans, i.e., 4487 times in (a), 4144 times in (b), 4869 times in (c), 4483 times in (d), 4371 times in (e), and 4788 times in (f).

## VII. Conclusions and Future Work

In this paper, we introduce a cost-efficient approach to refactor the latest version of FreeSurfer to support GPUs. To refactor such a large existing project that has tens of million of lines of source code, OpenACC directives are a more affordable approach, as an equivalent CUDA implementation requires significant programming efforts.

Experimental results demonstrate the usefulness and potential of using OpenACC directives to accelerate a large project like FreeSurfer on a GPU. An overlapped-fully-transfer scheme outperformed a copy-and-transfer scheme in reducing the side effects imposed by the CPU-to-GPU data transfer, achieving a speedup of 2.3× compared to the original CPU-based version. An average speedup of 1.2× for the entire GPU-accelerated module was obtained by the overlapped-fully-transfer scheme. Our approach is sustainable because it can be easily extended to other time-consuming functions and modules of FreeSurfer.

Moreover, in the process of implementing GPU support for FreeSurfer, we also gained knowledge that other users may use to smoothly utilize OpenACC to accelerate large projects. We also believe that the OpenACC group could use these learnings to improve the usability of OpenACC. The first is that we must avoid using `−ta=tesla:pinned` because this compiler option replaces all normal memory allocations with pinned memory allocations. Such a policy impairs the performance of a large project where not all the functions benefit from pinned memory. Instead, CUDA APIs to allocate and free pinned memory should be applied to only the necessary data. OpenACC will be more developer-friendly if they add such a fine-grained memory control feature to their `data` directives. The second is that we must avoid using `float` type variables to accumulate the computation results of `double` values. Because the possible precision loss will be different in CPU and GPU codes. For a normal program, such a small difference may not cause big problems, but for a large project where many iterations exist, the accumulated difference results in different behaviors in CPU and GPU codes. Manually performing such a search-and-fix process is cumbersome so it would be helpful if the OpenACC framework could implement a functionality to handle the job automatically.

Future work includes (1) reordering and fusing kernels to achieve better effects in overlapping data transfer overhead with computation time, and (2) exploring the possibility of kernel-level parallelism for higher efficiency in GPU resource utilization.

## Acknowledgments

This study was supported in part by the Japan Society for the Promotion of Science KAKENHI under grants 20K21794, and “Program for Leading Graduate Schools” of the Ministry of Education, Culture, Sports, Science, and Technology, Japan.

Data used in preparation of this article were obtained from the Alzheimer’s Disease Neuroimaging Initiative (ADNI) database (adni.loni.usc.edu). As such, the investigators within the ADNI contributed to the design and implementation of ADNI and/or provided data but did not participate in analysis or writing of this report. A complete listing of ADNI investigators can be found at: http://adni.loni.usc.edu/wp-content/upload-s/how_to_apply/ADNIAcknowledgement_List.pdf.

